# Sex differences in patch-leaving foraging decisions in rats

**DOI:** 10.1101/2023.02.19.529135

**Authors:** Marissa Garcia, Sukriti Gupta, Andrew M. Wikenheiser

**Affiliations:** Department of Psychology, University of California, Los Angeles, Los Angeles, California 90095; Current address: Neurosciences Graduate Program, University of California, San Diego, San Diego, CA 92093; Brain Research Institute, University of California, Los Angeles, Los Angeles, California 90095

**Keywords:** Foraging, decision making, marginal value theorem, patch

## Abstract

The ubiquity, importance, and sophistication of foraging behavior makes it an ideal platform for studying naturalistic decision making in animals. We developed a spatial patch-foraging task for rats, in which subjects chose how long to remain in one foraging patch as the rate of food earnings steadily decreased. The cost of seeking out a new location was varied across sessions. The behavioral task was designed to mimic the structure of natural foraging problems, where distinct spatial locations are associated with different reward statistics, and decisions require navigation and movement through space. Male and female Long-Evans rats generally followed the predictions of theoretical models of foraging, albeit with a consistent tendency to persist with patches for too long compared to behavioral strategies that maximize food intake rate. The tendency to choose overly-long patch residence times was stronger in male rats. We also observed sex differences in locomotion as rats performed the task, but these differences in movement only partially accounted for the differences in patch residence durations observed between male and female rats. Together, these results suggest a nuanced relationship between movement, sex, and foraging decisions.

**Significance statement:** Foraging behavior offers a naturalistic way of studying temporal investment amongst different choice options, a translationally-important form of decision making. Previous laboratory investigations have relied on foraging tasks that require little movement from subjects, which could affect the strategies that animals use and the neural mechanisms that support them. We developed a spatial foraging task for rats. Behavior generally matched the predictions of theoretical models, although rats remained in patches for longer than prescribed. Male rats exhibited a stronger tendency to overharvest patches than female rats. Sex differences in movement did not account for sex differences in foraging. These data highlight the interplay between movement and decision making, and demonstrate the utility of spatial tasks for studies of foraging.

## Introduction

Decision making is often studied by examining how subjects choose from a set of simultaneously-available outcomes. However, for decisions that have a temporal component, it is often equally important to decide how long a chosen resource should be exploited before moving on to a new option (Hayden, 2018; Stephens, 2008). For example, when reading a journal article, glancing at the figures might provide a broad understanding of the results quickly. Reading the introduction, results, and discussion sections provides further nuance and context. Exhaustively reading the paper and all of its supplemental information provides still more detail, but at this point the information gained by spending additional time with the paper will have decreased substantially. Many decisions involve such a tradeoff between persisting with an option as returns decrease or moving on to a new option. Several neuropsychiatric disorders are associated with reduced sensitivity to diminishing returns and a tendency to pursue options beyond the point that benefits outweigh costs (Raio et al., 2022; Speers & Bilkey, 2023), suggesting that this class of decisions has translational importance.

Animals face a similar problem when deciding how long to remain in foraging patches (MacArthur & Pianka, 1966; Stephens & Krebs, 1986; Stephens et al., 2007). As a forager harvests food, the patch they currently occupy depletes, causing their rate of food intake to decrease over time. Eventually, it will be advantageous to seek out a replete patch elsewhere. However, because food is not available in the locations between patches, it may be costly to travel to a new patch. Sophisticated foragers should therefore balance the diminishing returns in their current location against the expected cost of travelling and the expected quality of patches elsewhere. The marginal value theorem (MVT) provides a means of computing the optimal patch residence time, which occurs when the rate of return from the current patch falls to the average rate of return over the environment as a whole (Charnov, 1976; Kolling & Akam, 2017; Stephens & Krebs, 1986). Previous studies have shown that the qualitative predictions of the MVT hold over a range of species, including humans (Constantino & Daw, 2015; Hall-McMaster et al., 2021; Harhen & Bornstein, 2023; Kolling et al., 2012), nonhuman primates (Barack et al., 2017; Blanchard & Hayden, 2015; Hayden et al., 2011; Turrin et al., 2017), and rodents (Kane et al., 2017, 2019, 2022; Killeen et al., 1981; López-Yépez et al., 2021).

For most species, foraging involves movement over a potentially sizeable home range (Rudebeck & Izquierdo, 2022; Russell et al., 2005). However, laboratory studies of foraging are frequently conducted in operant boxes or other situations where subjects’ movements are relatively constrained. This difference may be consequential; when foraging requires movement between distinct locations, the parts of the brain responsible for spatial processing effectively become a part of the foraging system (Gordon et al., 2021, 2022). This means that while the computational goal of efficient foraging may be the same across topologically-equivalent behavioral tasks, its neural underpinnings may well differ. For example, in a scenario where patches are distinct spatial locations, spatial learning processes might enable a forager to form location-reward associations that track the quality of individual patches. However, if the unique reward statistics that define different patch types are experienced within a single spatial context, location-reward associations may be unhelpful—or even counterproductive—for distinguishing patch types.

For these reasons, a foraging task that involves locomotion between distinct locations would be a valuable complement to existing approaches for studying patch foraging. Here, we examined patch-leaving decisions in male and female rats on a patch foraging task with spatially-distinct patches. The reward statistics assigned to each patch varied unpredictably with every behavioral session, requiring subjects to update their location-reward associations flexibly. Rats’ behavioral strategies were compared to strategies that maximized food earnings, according to numerical simulations of the task.

## Materials and methods

*Subjects.* Long-Evans rats (n = 10, five female) weighing 200-300 g and aged 10-12 weeks at the start of the experiment were purchased from Charles River. Rats were housed singly and maintained on a 12-hour standard light/dark cycle. During task performance, rats were food restricted. Rats were weighed daily and provided supplemental chow in addition to what they earned on the behavioral task to maintain their weight at >85% of their free-feeding level. We did not track levels of circulating sex hormones in either male or female rats. All procedures followed the NIH Guide for the Care and Use of Laboratory Animals and were approved by the Chancellor’s Animal Research Committee at the University of California, Los Angeles.

### Apparatus

The test apparatus consisted of two square-shaped open field arenas (77 cm) connected to one another by a corridor (25 × 77 cm). At each end of the corridor, 12 cm openings could be blocked by doors actuated by stepper motors. Large extra-maze cues were visible from the foraging patches, allowing rats to orient themselves in space and discriminate between foraging locations. An infrared video camera (OptiTrack Flex 3) mounted overhead interfaced with video tracking software (Blackrock Neuromotive) to measure rats’ location in real time at 30 Hz. Two 40-mg food pellet dispensers (Med Associates) were positioned above each foraging patch to ensure an even distribution of food pellets across the foraging arenas. The behavioral task was controlled by custom Matlab (Mathworks) code. Behavioral events and video tracking data were timestamped by an electrophysiology recording system (Blackrock Cereplex direct).

### Behavioral task

Rats were handled 10-15 minutes per day for a week before behavioral testing began to acclimate them to contact with experimenters. During the week of handling, rats were also habituated to the behavioral apparatus by allowing them to explore it for approximately 10 minutes per day with all doors open, and with sucrose pellets scattered randomly throughout the two foraging locations. Following handling and apparatus habituation, rats performed one 30- minute task session each day. At the beginning of each session rats were placed in the travel corridor with doors to both patches closed. When the programmed travel delay for that session elapsed, one of the two patch doors was randomly selected to open, and the rat was free to enter the patch and begin foraging. While rats occupied a foraging patch, food pellets (40 mg, 100% sucrose; Bioserv) were delivered following the time-based reward schedule assigned to the patch in that session. Rats could choose to abandon their current foraging patch at any time by entering the travel corridor and approaching within 10 cm of the door to the opposite patch; this caused the door to the previous patch to close, confining the rat to the travel corridor for the travel time assigned to that session. When the travel time elapsed, the door to the next patch opened. Rats were forced to visit patches in an alternation pattern, though there was no minimum amount of time rats were required to remain within a patch. Food pellets were not delivered to either patch while rats were in the travel corridor.

### Reward schedules

Patch reward schedules were parameterized by an initial inter-pellet interval (which set the rate of reward when rats first entered the patch), and an exponential time constant (which determined how quickly inter-pellet interval increased while rats remained within the patch). These parameters defined food pellet delivery rate as a function of patch residence duration. The precise time of individual pellet deliveries was determined by drawing a latency from a Gaussian distribution with mean equivalent to the reciprocal of the current rate of reward and a standard deviation of four seconds. In this way, the cumulative number of pellets earned for a given residence time matched the functions plotted in figure 1b on average, but the precise timing of pellet deliveries was unpredictable on successive visits to the same patch. Three reward schedules were generated by fixing the initial inter-pellet interval to 15 s and setting the exponential time constant that determined how quickly inter-pellet interval increased over time to 100 s, 300 s, and 600 s, corresponding to fast, medium, and slow patch depletion rates (fig. 1c). Additional reward schedules were created by fixing the exponential time constant that determined how quickly inter-pellet interval increased over time to 300 s and varying the initial inter-pellet interval to create high, medium, and low initial reward rates (fig. 1c). For half of the rats, the initial inter-pellet intervals were 10, 15, and 20 seconds for the high, medium, and low initial reward rate schedules. For the remaining half of the subjects, the initial inter-pellet intervals were 12, 15, and 18 seconds for the high, medium, and low initial reward rate schedules. The effect of varying initial reward rate was qualitatively similar for these slight differences in initial reward rate, so data are combined in figure 3. However, we used the actual initial inter-pellet intervals as predictors in the mixed-effects models and numerical simulations of the task described below.

**Figure 1.**
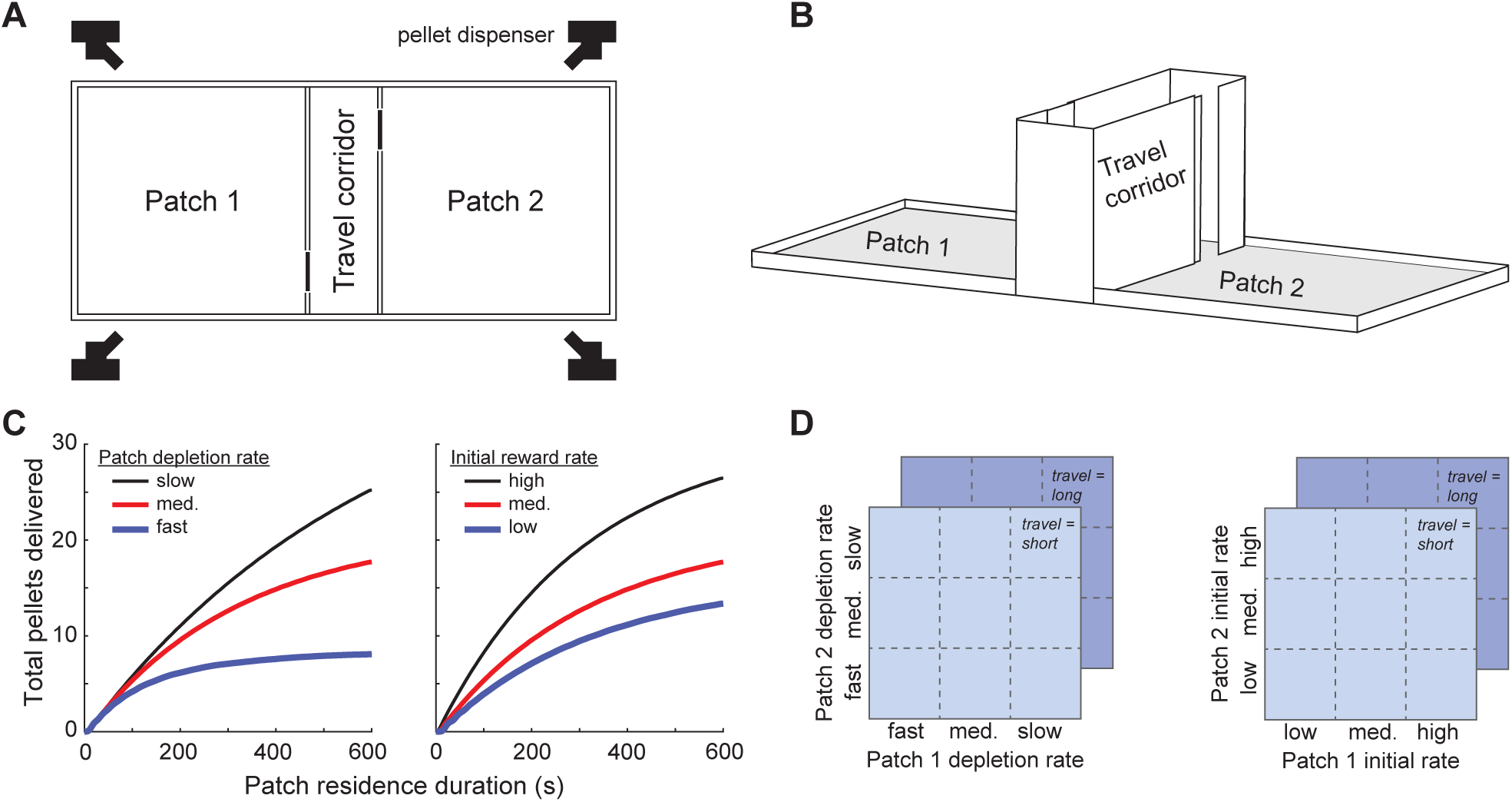
A spatial patch foraging task for rats. (A) The behavioral apparatus consisted of two open field foraging patches connected by a rectangular corridor. Doors at each end of the corridor controlled access to different parts of the apparatus. Food pellet dispensers mounted above each patch distributed food evenly across both foraging patches. (B) Three-dimensional view of apparatus, depicting the door leading to patch two in the open position and the door leading to patch one in the closed position. (C) Reward schedules determined how many food pellets were delivered on average while rats remained within a patch. Three schedules were created by varying the rate at which patches depleted (left). Three additional reward schedules were created by holding patch depletion rate constant but varying the initial rate of food delivery (right). (D) Rats experienced combinations of the reward schedules depicted in (C) along with two different travel times in pseudorandom order.

### Testing sequence

After handling, rats performed five training sessions with short travel time, medium depletion rate, and medium initial rate. By the fifth training session all rats were reliably switching between patches multiple times within each sessions and were deemed ready to begin the task sequence. Behavioral sessions were constructed by assigning reward schedules to each of the patches, and choosing a travel time; these assignments remained fixed for the duration of the session. We tested all assignments of the depletion rate and initial reward rate schedules to the physical patch locations with both long (60 s) and short (0 s) travel times (fig. 2c). Rats progressed through the sequence of sessions in a block-wise fashion, with half of rats first completing the depletion rate sessions and the other half first completing the initial rate sessions. Within each block, the order of sessions was randomized. This design resulted in the depletion rate and initial rate blocks each comprising 18 sessions, for a total of 36 sessions per rat, and 360 sessions for the entire cohort of 10 rats. Two sessions from the depletion rate block (from two different rats) were discarded due to errors in saving data that were not discovered until after the experiment was complete. Additionally, one rat was inadvertently tested on the same set of session parameters from the initial rate block twice; there was no principled reason to prefer one of these sessions over the other, so both were included in the final data set. Together, a total of 359 sessions (181 initial rate manipulation sessions, 178 depletion rate manipulation sessions) were included for analysis.

**Figure 2.**
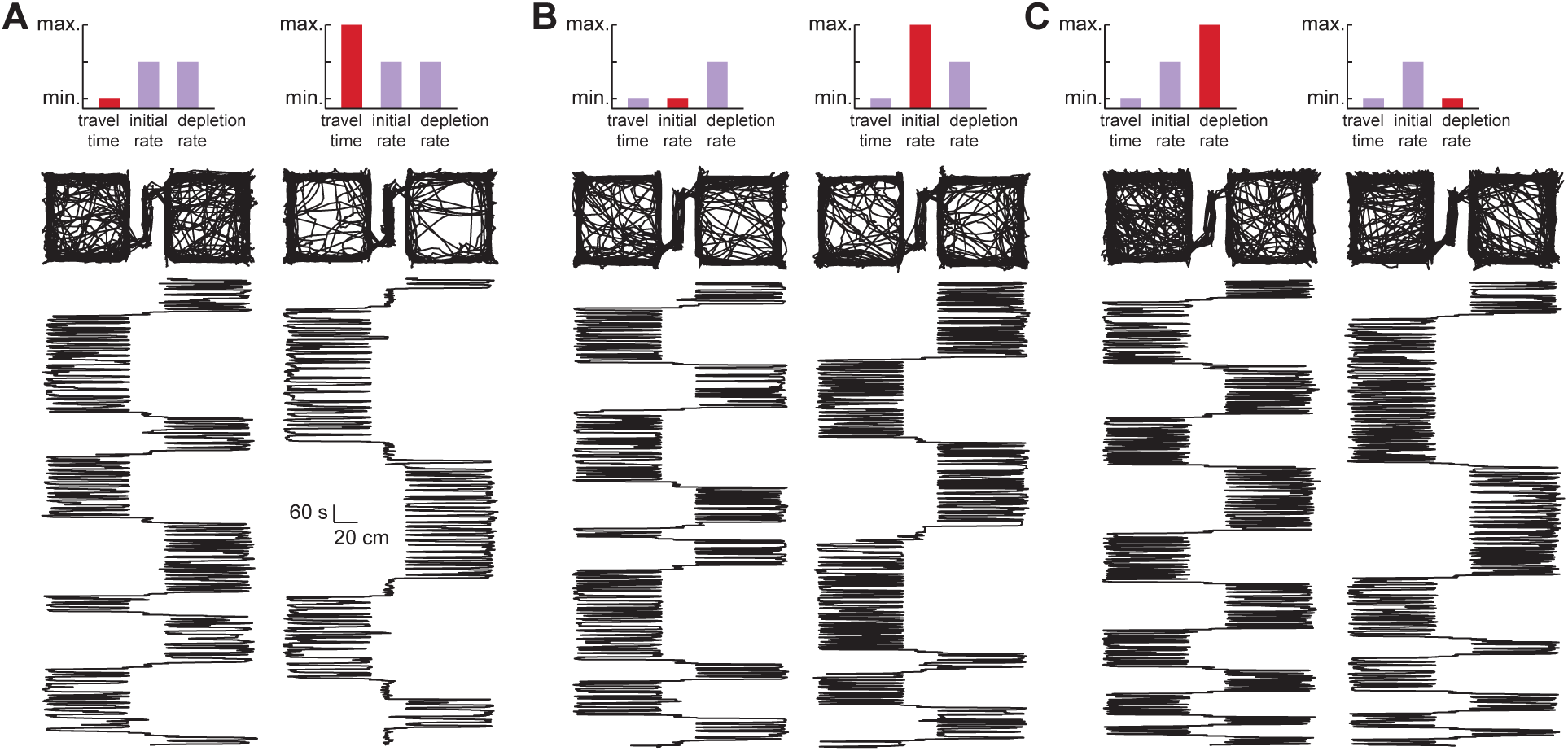
Example behavioral sessions. Each panel displays position-tracking data for an example session, in a top-down view (top row), and with the rat’s x-coordinate position plotted against session time increasing along the y-axis (bottom row), such that session start is at the bottom of the figure. Pairs of sessions that differed in a single task parameter were selected to show how patch residence time was affected by differences in travel time (A), initial reward rate (B), and patch depletion rate (C).

### Data analysis

All data analyses were conducted in Matlab (2021a). Patch residence time was computed for each patch in each session by taking the median duration of all visits to the patch. The start time of a patch visit was taken to be the moment when the rat’s center of mass crossed from the travel corridor to the foraging patch. The end time of visits was taken to be the moment that rats approached the door to the opposite patch and triggered the travel time phase of the task. Visits to a patch that were interrupted by the end of the session were not included in the calculation of patch residence time because there was no way of knowing how much longer the rat would have remained in the patch.

All models throughout the paper were linear mixed effects models that were fit using the Matlab function *fitlme*. These models allow us to quantify how a dependent variable of interest is modulated by one or more predictors. Importantly, mixed-effects models allow for subject identify to be included as a random effect, which accounts for the dependence in data arising from repeated measurements within individual rats (Yu et al., 2022). Thus, for all models reported throughout the paper, subject identity was included as a random effect. Continuous variables were z-scored so reported coefficient estimates are standardized. For the model assessing how task parameters and subject factors affected patch residence time (table 1), fixed effects were initial patch reward rate, patch depletion rate time constant, and travel time (coded as continuous variables), as well as sex and patch location (coded as categorical variables; females and left patch coded as “one”). Subject identity was included as a random effect and coded as a categorical variable. We also tested a version of the model that included the predictors described above, as well as every pairwise interaction between predictors. However, none of the interaction terms were significant, so we excluded interactions from the model reported here to maximize the accuracy of coefficient estimates.

**Table 1.** Regression analysis of patch residence duration.

| Effect | Estimate | SE | 95% CI |  | *p* |
| --- | --- | --- | --- | --- | --- |
|  |  |  | *LL* | *UL* |  |
| Intercept | -3.19×10<sup>-16</sup> | 0.03 | -0.06 | 0.06 | 0.99 |
| Depletion rate, current patch | -0.20 | 0.03 | -0.24 | -0.13 | 4.00×10<sup>-10</sup> |
| Depletion rate, opposite patch | 0.10 | 0.03 | 0.03 | 0.14 | 2.90×10<sup>-3</sup> |
| Initial rate, current patch | 0.13 | 0.02 | 0.08 | 0.19 | 5.05×10<sup>-6</sup> |
| Initial rate, opposite patch | -0.09 | 0.03 | -0.14 | -0.03 | 4.44×10<sup>-3</sup> |
| Travel time | 0.55 | 0.02 | 0.49 | 0.60 | 2.30×10<sup>-63</sup> |
| Sex ("female" coded as 1) | -0.13 | 0.03 | -0.18 | -0.07 | 1.53×10<sup>-5</sup> |
| Side ("left" coded as 1) | -0.03 | 0.03 | -0.10 | 0.01 | 0.14 |

LL, lower limit of confidence interval; UL, upper limit of confidence interval; SE, standard error

For the model examining the influence of running speed on patch residence duration, fixed effects were mean running speed within the patch (a continuous variable), sex (a categorical variable; females coded as “one”), and their interaction. Because the interaction was significant in this case, we retained it for the model reported in the text. Subject identity was included as a categorical random effect.

### Determining rate-maximizing patch residence times

A behavioral strategy on the task comprised the set of patch one and patch two residence durations. We considered only fixed, deterministic strategies as there was no way for rats to predict the stochasticity of the reward schedules, and therefore no way in which choosing a different residence duration on successive visits to the same patch could produce a better outcome than consistently applying the correct deterministic strategy. For each pair of residence durations (ranging from 0 – 600 s in one second increments) we used numerical simulations to compute the associated food pellet earnings. Simulations began with the session time set to zero seconds; as in the actual behavioral task, the first event to occur was the programmed travel duration, so session time was incremented by that amount. Next, one patch was randomly selected to be the first available to the simulated agent. For each simulated patch visit, food pellet delivery times were calculated according to the reward schedule assigned to the patch as described above, and any pellets delivered before the agent’s residence duration for that patch elapsed were added to the agent’s total earnings. The session time was then incremented by the patch residence duration and the travel time assigned to that session, and the agent then moved on to the second patch. The programmed travel delay for the short travel time was zero seconds; however, rats were not able to transit the travel corridor instantaneously. The average transit time across all short travel time sessions was approximately 5 s, so for simulations of sessions with a short travel time the session time was incremented by 5 s when rats switched patches. Simulations continued in this way until the session time reached 30 minutes, the duration of actual behavioral sessions. Using this method, we computed the rate of reward over the strategy space of patch one and patch two residence times (fig 5a). Each pair of residence durations was simulated 500 times for each session type, and earnings were averaged across simulations. The resulting rate maps were smoothed by 1 second in each direction, and the peak pixel was taken to be the rate-maximizing behavioral strategy for that session type. For the figure showing behavior and model-predicted reward rate (fig. 5a), we included data from symmetrical sessions (i.e. sessions involving the same the same two reward schedules and differing only in which schedule was assigned to the left or right patch) when possible, as there was no significant effect of position within the apparatus.

### Journeys & movement statistics

Artifacts in video tracking data (i.e. large jumps in position) were removed and corrected by interpolation. Following previously-described approaches, we segmented position-tracking data into continuous bouts of movement (journeys) and periods with little or no sustained movement. Periods when rats were confined to the travel corridor were not included in this analysis as movement was limited by the confines of the corridor. The onset of a journey was defined as the moment running speed exceeded 5 cm/s, and the journey terminated when running speed fell below 5 cm/s. The distance travelled on each journey detected in this way was computed by summing the rat’s displacement between successive tracking points; very short journeys (total distance < 20 cm) were excluded from analysis. Features of the remaining journeys (average running speed, distance travelled, and distance to center of patch) were computed. We categorized journeys as center crossing if the rat approached within 15 cm of the center of the arena at any point on the journey. Independent of journeys, we computed the total distance travelled in each session while foraging in the patches (total path length) by summing the distance between successive positions, excluding intervals when the rat was in the travel corridor.

To examine the distribution of behavior over space we divided the apparatus into 1-cm bins and determined how long rats spent at each location in each session. Occupancy at each location was averaged across sessions separately for male and female rats (fig. 6E). We used the average male and female occupancy to compute the occupancy difference index for each location on the apparatus (fig. 6F). Occupancy difference index was defined as the difference of female and male average occupancy divided by the sum of male and female average occupancy for each location; in this way, values near zero indicated relatively equivalent male and female occupancy, while positive (negative) values indicated proportionately greater time spent at a location by female (male) rats.

## Results

### A laboratory test of patch-leaving decisions

We tested rats (n = 10; five female) on a foraging task designed to mimic the structure of natural patch-leaving decisions. The test apparatus (fig. 1a-b) consisted of two open-field foraging patches connected by a corridor. Computer-controlled doors at each end of the corridor controlled access to the patches and dispensers mounted above each patch allowed food pellets to be delivered. Rats began each session in the travel corridor, with doors to both patches closed. Next, one randomly-selected door opened, allowing access one foraging patch. On entering the accessible patch, food pellet delivery began and rats were free to forage for as long as they cared to. To switch between patches, rats entered the travel corridor and approached the closed door to the opposite patch. This caused the door to the recently-exited patch to close, and the rat was then confined to the travel corridor for either a long (60 s) or short (0 s) travel time delay to simulate time spent moving between patches. Food was never available in the travel corridor, so switching patches was costly in terms of lost opportunity to collect food pellets. When the travel time elapsed, the door to the unvisited patch opened, and the rat was free to enter and begin foraging.

Food pellets were delivered at unpredictable intervals according to a time-dependent schedule (fig. 1c). In all cases, patches depleted as rats remained in them, with the rate at which food pellets were delivered decreasing over time. For some schedules the rate of reward began at the same initial value and patch depletion occurred more or less quickly (fig. 1c, left); in other cases, the rate at which reward decreased was held constant but the initial reward rates differed (fig. 1c, right). Each time that rats re-entered a patch the reward schedule reset to its maximum rate. For each behavioral session, one gain function was assigned to each patch and one travel time was selected. The combinations of task parameters we tested comprised 36 unique session types that each rat experienced once in pseudo-random order (fig. 1d).

### Sex differences in patch-leaving decisions

Figure two shows position-tracking data for example sessions, with the rat’s x-coordinate position plotted over time (start of session at the bottom of plot), a view that makes it easy to visualize how much time rats spent in the left and right patches. The pairs of example sessions differed in travel time (fig. 2a), initial reward rate (fig. 2b), or reward depletion rate (fig. 2c), with otherwise identical task parameters. When travel time was short, the rat switched between patches frequently (fig. 2a, left); however, when travel time was long, the rat remained in patches for a longer duration on each visit (fig. 2a, right), consistent with the predictions of foraging models. Rats similarly spent longer in patches that had a higher initial reward rate (fig. 2b, right) and a slower reward depletion rate (fig. 2c, right).

We computed patch residence duration as the median duration of all visits to a patch within a behavioral session. Patch residence duration increased with decreasing reward depletion rates and with increasing initial reward rates (fig. 3a). Rats were also sensitive to travel time, with longer travel times resulting in increased residence durations for all reward schedules (fig. 3a). Further, the amount of time that rats spent in one patch was influenced both by the local reward schedule, and by the reward schedule of the opposite patch (fig. 3b). When the opposite patch reward schedule was more advantageous than that of the current patch (i.e. higher initial rate or slower depletion rate), rats reduced the amount of time they spent in the current patch relative to situations when both patches had identical schedules. Similarly, if the opposite patch had a less favorable reward schedule than the current patch, rats increased their allocation of time to the current patch. These results indicate that rats integrated multiple independent sources of cost and benefit information in a manner broadly consistent with the predictions of theoretical foraging models.

**Figure 3.**
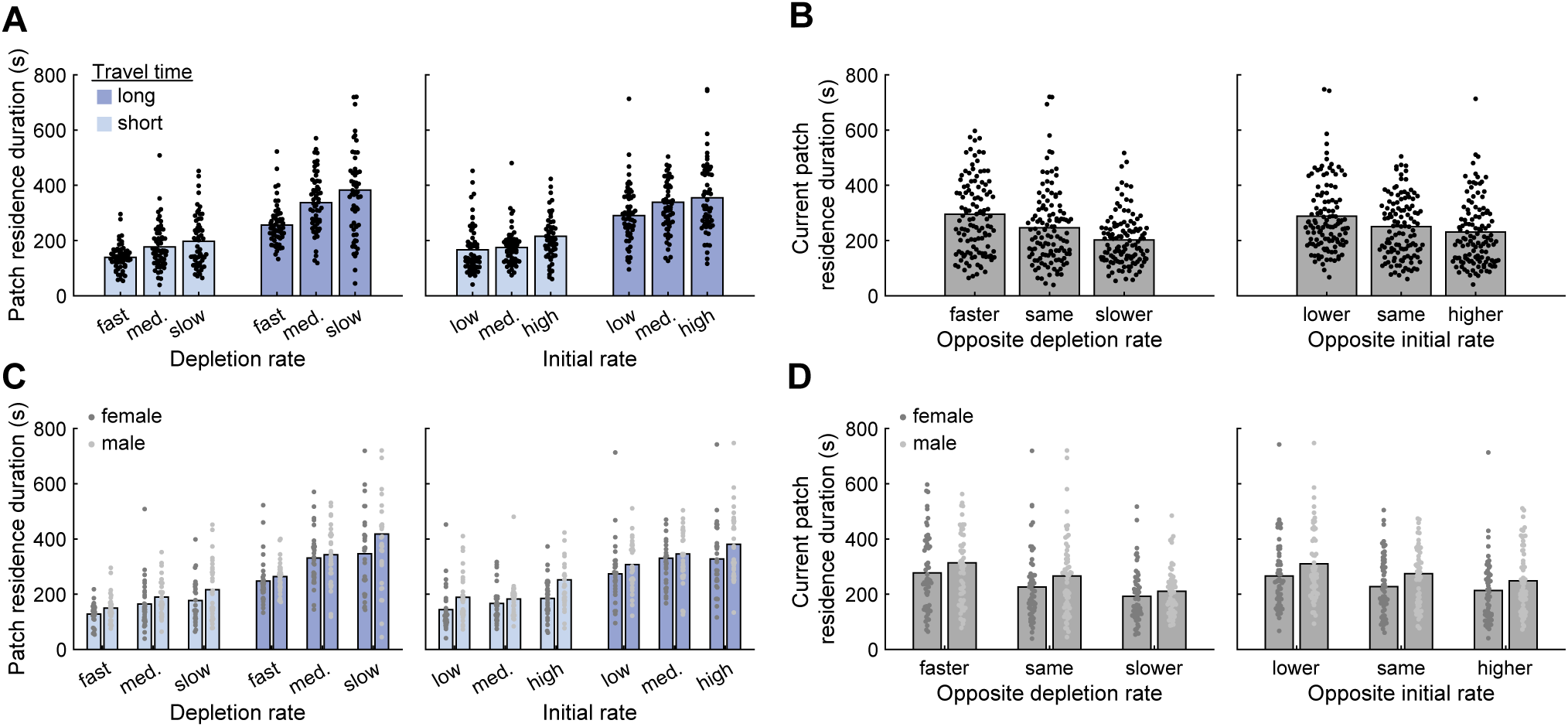
Sex differences in foraging. (A) Patch residence duration is aggregated by depletion rate (left), initial rate (right), and travel time (both plots). Each point depicts patch residence duration for one rat in one patch in one behavioral session; bars indicate group means. (B) Patch residence duration is aggregated based on whether the other patch in the session had more favorable reward statistics (i.e. slower patch depletion rate or higher initial rate), less favorable reward statistics (i.e. faster patch depletion rate or lower initial reward rate), or identical reward statistics. Each point depicts patch residence duration for one rat in one patch in one behavioral session; bars indicate group means. (C) Same data presentation as in (A), except that data are separated by sex. (D) Same data presentation as in (B), except that data are separated by sex.

Interestingly, we observed clear sex differences in patch residence times, with female rats systematically exiting patches earlier than male rats. This difference was consistent across conditions: female rats showed numerically lower residence times compared to male rats in every session type that we tested (fig. 3c). Female and male rats were similarly sensitive to local patch and opposite patch reward schedules (fig. 3d).

We used linear mixed-effect models to quantify how task parameters and subject-specific factors were related to patch residence duration. Fixed effects in the model included sex, reward schedule parameters (initial reward rate, patch depletion rate) for both the currently-occupied and opposite patches, travel time, and the physical location (left or right) of the patch within the testing room. Subject identity was included as a random effect. Travel time and reward schedule parameters exerted significant effects on patch residence duration in the directions predicted by classical models of foraging behavior (table 1). Sex was a significant predictor of residence duration, consistent with the results shown in figure three. The physical location of patches within the apparatus did not significantly affect residence duration. Though we initially tested a version of the model that included all two-way interactions between predictors, none were found to be significant, so interaction terms were not included in the final model.

We also tested whether parameters from previous sessions influenced rats’ behavior. We fit a mixed-effects model to examine how patch residence time was influenced by travel time, depletion rate, and initial reward rate, both for the current session parameters and those of the five preceding sessions. As with our previous analysis, this model included sex and side as fixed effects and subject identity as a random effect.

Standardized coefficient estimates are plotted in figure 4. Session 0 corresponds to the current session, and negative values for session number denote previous sessions. The current session travel time (β = 0.55, p = 2.02 × 10^-54^), patch depletion rate (β_current patch_ = -0.19, p = 7.92 × 10^-10^; β_opposite patch_ = 0.09, p = 0.003), and initial reward rate (β_current patch_ = 0.15, p = 3.54 × 10^-6^; β_opposite patch_ = -0.07, p = 0.02) all significantly influenced patch residence time. The travel time from the previous session also significantly influenced patch residence time (β = 0.11, p = 0.001). The effects of patch depletion rate and initial reward rate were limited to current session values only, with no significant influence of previous values for these variables. These results suggest that travel time had a lingering effect on behavior, indicating that rats integrated their estimates of some environmental statistics over a timescale longer than a single session.

**Figure 4.**
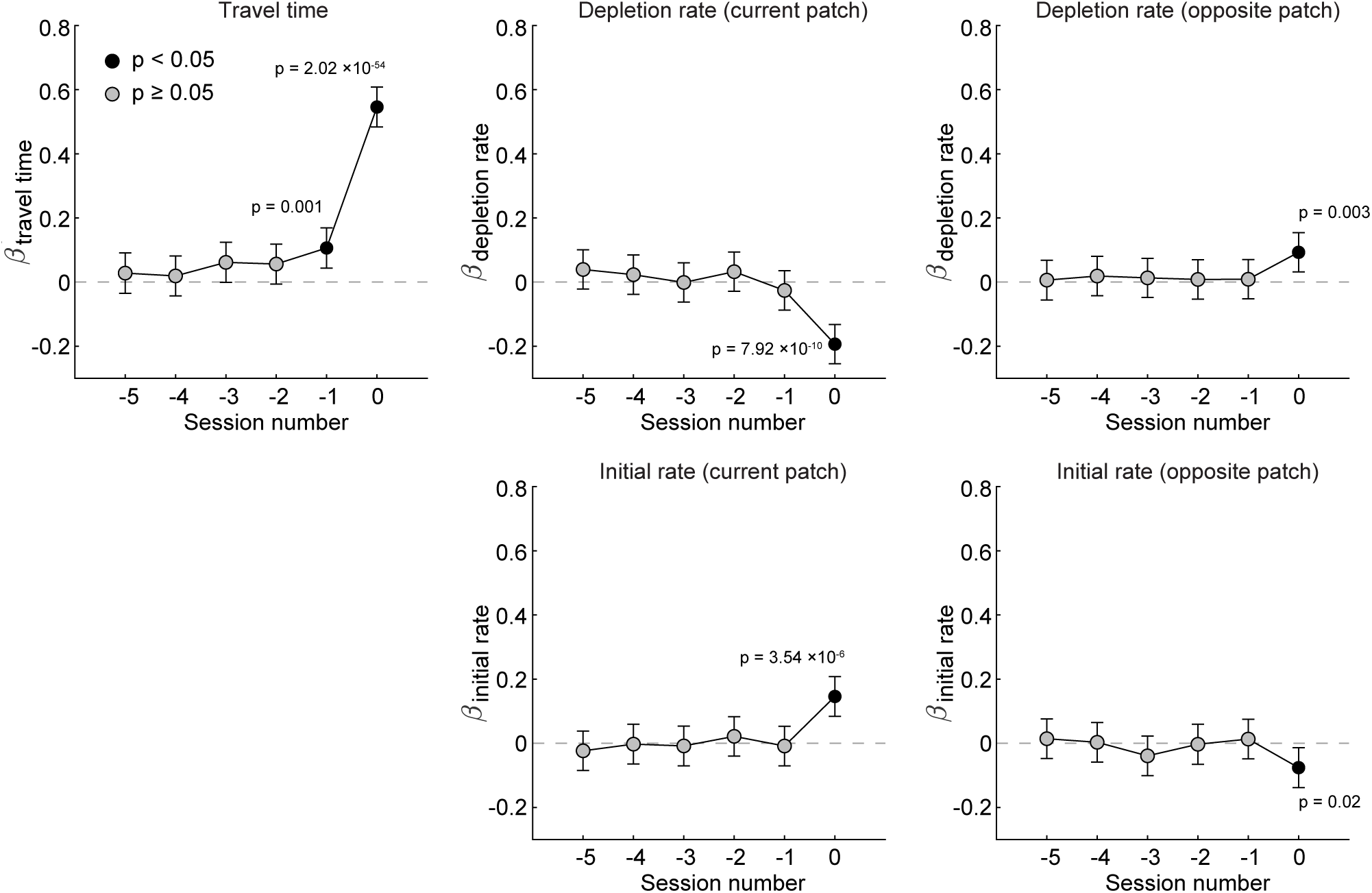
Session history effects. We assessed whether the travel time, reward depletion rate, and initial reward rate of previous sessions affected patch residence time, for both rats’ current patch, and the opposite, unoccupied patch. Consistent with earlier analyses, the present values of all of these variables (i.e. session number 0) significantly affected patch residence time. In addition, the travel time of the previous session significantly affected residence time. Values of depletion rate and initial rate from previous sessions did not significantly modulate rats’ behavior.

### Rats overharvested patches

We used numerical simulations to determine the patch residence durations that resulted in the greatest food earnings for each session type. Assuming a fixed residence duration for each visit to a patch, we computed the rate of reward associated with all possible combinations of patch one and patch two residence durations. In figure 5a, lighter and darker colors indicate higher or lower average reward rates across combinations of patch residence durations in three example session types. Observed residence durations for male and female rats are also plotted. Rats’ behavior generally tracked regions of strategy space associated with high rates of reward. However, both male and female rats systematically chose patch residence durations longer than the rate-maximizing residence durations determined by simulation. Across all sessions, rats’ chosen residence durations were strongly correlated with (r = 0.55, p = 3.54 × 10^-57^, n = 718 patch residence times, Pearson’s correlation; fig. 5b), but systematically greater than, rate-maximizing residence durations (mean optimal patch residence time = 153 s; mean observed patch residence time = 252 s; β_observed_ = 0.81, p = 1.00 × 10^-29^; linear mixed-effects model). Though both male and female rats overharvested patches, female rats overharvested significantly less (β_female_ = -0.30, p = 5.62 × 10^-5^; linear mixed-effects model; fig 5c). By leaving patches earlier than male rats, female rats more closely approximated the rate-maximizing strategy; consistent with this, female rats earned slightly more food pellets per session compared to male rats (55.1 vs. 51.1 pellets; β_female_ = 0.24, p = 0.02; linear mixed-effects model).

**Figure 5.**
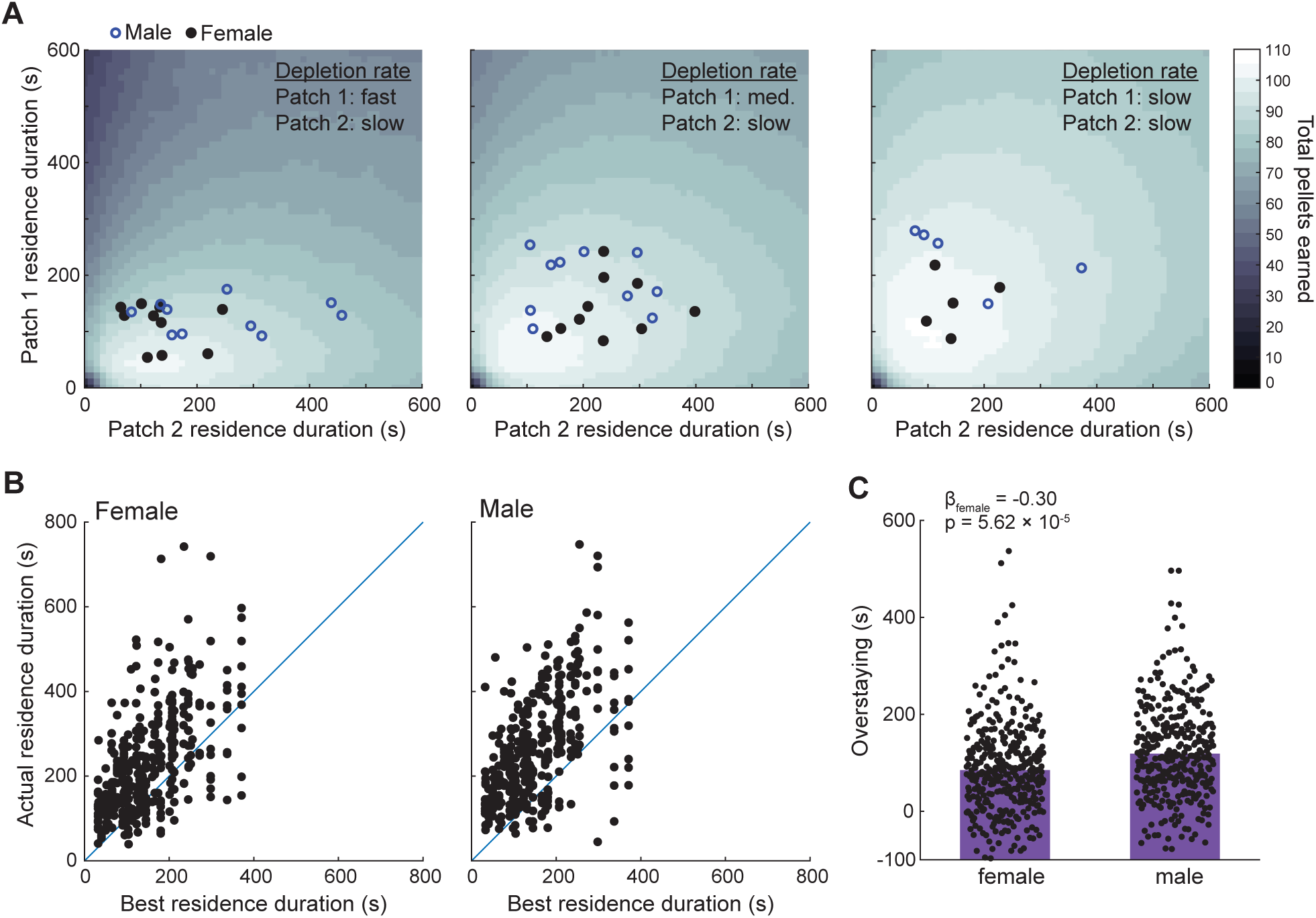
Rats overharvested patches. (A) In each panel, heat maps depict food earnings resulting from each behavioral strategy (i.e. each pair of patch residence durations) as determined by simulations. Results for three session types are shown, with patch one depletion rate ranging from fast to slow, and all other task parameters fixed (travel time = 60 s; patch 2 depletion rate = slow). Data points depict observed rat behavior for sessions with matching parameters; each point represents behavior from one rat in one session. (B) Observed patch residence duration is plotted against the patch residence duration that earned the greatest amount of food in simulations; each point depicts results from one rat in one patch in one behavioral session. (C) Overstaying (the difference between the simulation-determined best residence duration and the observed residence duration) is aggregated by sex. Each point depicts one patch in one behavioral session. Blue crosses depict the mean for each distribution. Positive values indicate that residence durations exceeded the best durations determined by simulation. Overstaying was significantly greater in male rats.

One possible cause of overharvesting is that rats earned enough food to reach satiety within sessions and were therefore under less pressure to choose patch residence times that maximized food intake. Several lines of evidence argue against this idea. First, the maximum amount of food any rat ever earned on the task (101 pellets, 4.5 grams) falls short of the amount of food rats need to maintain weight from day to day. Average food earnings per session (53.1 pellets, 2.4 g) were lower still, suggesting that rats were unlikely to have reach satiety from task earnings alone. Second, the satiety account predicts that overstaying should be more pronounced in richer sessions where rats earned many food pellets due to short travel times and favorable patch depletion statistics. However, we observed a mild but significant negative correlation between the number of food pellets rats earned and magnitude of overstaying (r = -0.19, p = 1.47 × 10^-7^; Pearson’s correlation), indicating that overharvesting was more prevalent in sessions where rats earned fewer pellets. Finally, female rats are smaller on average than male rats, and should therefore reach satiety more quickly; if overharvesting was driven by satiety it should be more pronounced in female rats. Instead, we observed the opposite, with female rats showing less overharvesting than male rats.

### Sex differences in movement patterns do not fully explain sex differences in foraging

Previous work has shown that male and female rats move through and interact with open spaces differently (e.g. Bishnoi et al., 2021; Hyde & Jerussi, 1983; Osterlund Oltmanns et al., 2021; Stöhr et al., 1998). Consistent with this, we observed differences between male and female rats in several movement properties. Following previous approaches (Drai et al., 2000; Eilam & Golani, 1989), we segmented position-tracking data into discrete journeys—bouts of continuous locomotion bounded by periods of little or no movement—and computed average journey properties for each behavioral session. Female rats traveled further on average for each journey (β_female_ = 0.81, p = 0.03; linear mixed-effects model; fig 6b), though there was no significant difference in their average journey movement speed (β_female_ = 0.75, p = 0.08; linear mixed-effects model; fig 6a). There was no difference in path length, the total distance rats travelled over the course of each session (β_female_ = 0.69, p = 0.11; linear mixed-effects model; fig 6c), though female rats were significantly more likely to make journeys that crossed through the center of the foraging patches (β_female_ = 0.57, p = 0.04; linear mixed-effects model; fig 6d). Indeed, examining time spent at each location across the apparatus by male and female rats revealed a tendency for male rats to spend relatively greater amounts of time along the perimeter of the foraging patches, and female rats to spend relatively greater time near the center of patches (fig. 6e-f).

**Figure 6.**
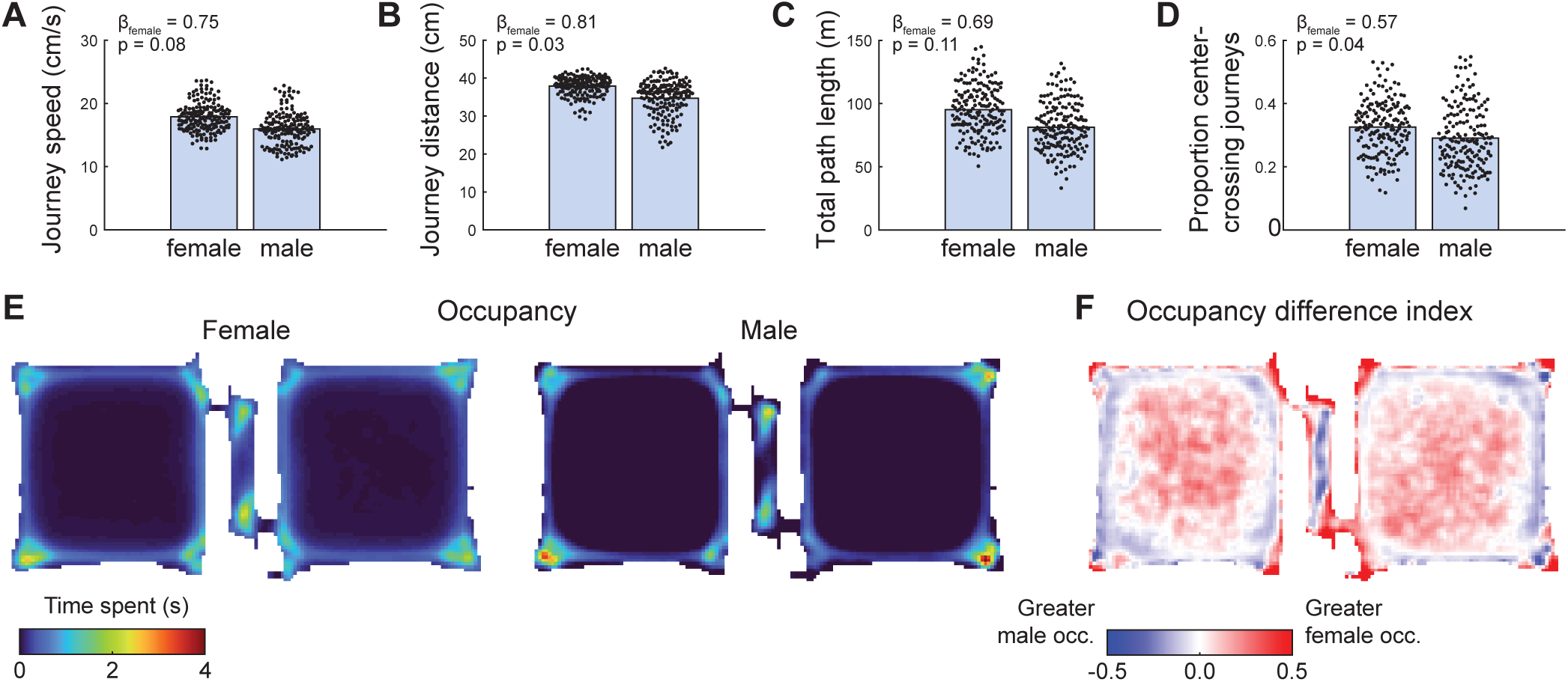
Sex differences in movement. (A) Average movement speed during journeys—discrete bouts of movement—was greater for female rats. Each data point shows average journey speed for one session; bars depict group means. (B) The distance travelled per journey was greater for female rats. Each data point shows average journey distance for one session; bars depict group means. (C) Total distance travelled per session (computed without segmenting data into discrete journeys) was greater for female rats. Each data point shows total distance travelled for one session; bars depict group means. (D) A fraction of female rat’s trajectories crossed through the center of the open field foraging patches. Each data point shows the fraction of total journeys within one session that passed within 15 cm of the patch’s center. (E) Heat maps show the amount of time female (left) and male (right) rats spent at different locations on the apparatus, averaged over all subjects and all sessions. (F) Heat map shows the occupancy difference index, the relative difference in time spent at each location between male and female rats. Redder colors denote locations where females spent more time on average than males, bluer colors indicate locations where males spent a greater amount of time on average than females. White denotes similar occupancy between males and females.

We wondered whether basal differences in movement could account for the sex differences in foraging decisions we observed. For instance, if rats moved randomly within patches until they encountered the door to the travel corridor and exited the patch, rats with a higher average running speed would reach doors more quickly and therefore exit patches earlier. This account predicts that variation in running speed should systematically affect patch residence duration, with faster running speeds resulting in shorter residence durations.

For individual visits to patches across all subjects and sessions (n = 2464 patch visits), we computed rats’ mean running speed while in the patch and fit a linear model to assess how well running speed, sex, and their interaction predicted patch residence duration. The model revealed significant effects of running speed (β_speed_ = -0.11; p = 1.34 × 10^-30^), sex (β_female_ = -0.91; p = 8.29 × 10^-22^), and the interaction of running speed and sex (β_speed*female_ = 0.10; p = 2.35 × 10^-17^). These results indicate that while running speed and sex both influenced how long rats remained in patches, the effect of sex was substantially larger. However, the significant interaction term indicates that the nature of the relationship between running speed and residence duration differed for males and females. Indeed, running speed and residence duration were negatively correlated in male rats (r = -0.31; p = 1.25 × 10^-26^; Pearson’s correlation), but these variables were not related in female rats (r = -0.03; p = 0.36; Pearson’s correlation). These findings suggest a nuanced relationship between running speed, sex, and patch-leaving decisions: male rats that moved more quickly also tended to leave patches earlier, but the fact that female rats ran faster on average than male rats does not fully account for their tendency to abandon patches earlier.

## Discussion

Foraging is a naturally motivated form of decision making that is essential for survival in most species (Calhoun & Hayden, 2015; Hayden, 2018; Stephens & Krebs, 1986; Stephens, 2008). Understanding the neural basis of foraging decisions holds great potential for understanding decision making generally, as it have been argued that the evolutionarily-ancient mechanisms that support foraging decisions may have been repurposed to support other types of decision making and cognition (Hills et al., 2008; Rudebeck & Izquierdo, 2022; Todd et al., 2012; Todd & Hills, 2020). Further, in recent years there has been growing recognition that studying natural behaviors like foraging may be an especially productive way to understand brain function (Cisek & Hayden, 2021; Gomez-Marin & Ghazanfar, 2019; Hein et al., 2020; Krakauer et al., 2017; Miller et al., 2022; Yoo et al., 2021). Testing decision making in a way that mimics the natural structure of foraging decisions increases the ethological validity of the paradigm by tailoring the relevant sensorimotor features of the task to match what animals would typically experience in natural settings (Juavinett et al., 2018). For all of these reasons, well-validated laboratory tests of foraging behavior are a valuable tool for understanding the neurobiology of decision making.

We tested male and female rats on a spatial patch-foraging task. Rats learned the local reward statistics assigned to spatially-distinct foraging patches and the travel time penalty associated with switching between them. Using this knowledge, rats adjusted the amount of time they allocated to each foraging location, paralleling—though not perfectly matching—the patch residence durations that maximized food earnings. These results show that the task elicited flexible, context-sensitive foraging behavior from rats. This work highlights the promise of spatial tasks as a means of studying the neurobiology of foraging decisions.

### Overstaying and patch foraging

As in previous studies of patch foraging involving a range of species, rats in our study consistently chose residence times longer than those predicted by food intake rate maximization models (Cash-Padgett & Hayden, 2020; Cassini et al., 1990; Constantino & Daw, 2015; Hayden et al., 2011; Kane et al., 2019; Nonacs, 2001). Many possible explanations have been proposed for this overharvesting phenomenon, and it is probably driven by a combination of different factors (Kendall & Wikenheiser, 2022). Perhaps the most general explanation—and one not exclusive of various other causes—is that rats were optimizing more than food earnings. In most scenarios foragers must consider many other factors while searching for food, including predation risk (Choi & Kim, 2010; Ydenberg & Dill, 1986), potential competitive or affiliative interactions with conspecifics (Li et al., 2012; Nagy et al., 2020; Whishaw, 1988; Whishaw & Tomie, 1988), energy expenditures (Hart et al., 2017), and opportunities for learning about aspects of their surroundings (Harhen & Bornstein, 2023; Wang & Hayden, 2021). Foraging strategies that balance these and other factors might depart from the predictions of rate maximization but nevertheless perform well in real-world scenarios. Future experiments that manipulate food availability alongside important non-food factors will be important for understanding the behavioral and neural mechanisms of the complex multi-objective optimization problem that natural foraging entails.

### Sex differences in foraging decisions

Female rats left patches earlier than male rats across all task conditions (fig. 3), and our regression analysis indicated that sex differences in running speed accounted for only a small portion of this effect. This suggests that a yet-unidentified difference between male and female rats drives divergent patch-leaving decisions. Gonadal hormones—acting over the course of development or on a shorter timescale—are a clear candidate mechanism for orchestrating sex differences in decision making. Indeed, a recent study of intertemporal choice in rats (Hernandez et al., 2020) reported robust sex differences that were controlled by circulating testosterone levels. When choosing between a small amount of food delivered quickly and a larger amount of food available after a delay, male rats preferred the “larger-later” option, while female rats opted for the “smaller-sooner” selection. Reducing testosterone levels in male rats shifted their preference toward the smaller-sooner choice, while reducing ovarian hormones in female rats did not alter their preferences. Interestingly, previous work has found that temporal preferences measured on intertemporal choice tasks are predictive of patch-leaving decisions (Kane et al., 2019), with a tendency to favor immediate over delayed rewards in the intertemporal scenario corresponding to overharvesting patches in the foraging scenario. Integrating these two lines of work, one might predict that female rats would be especially prone to overstaying—the opposite of what we observed. However the different strains of rats used here (Long-Evans) and in Hernandez et al.’s (2020) work (Fisher-Brown Norway hybrid) could also impact the relationship between sex and intertemporal choice. More work investigating how sex, strain, and hormones interact on foraging and intertemporal choice tasks would clearly be fruitful.

Our results add to the growing list of sex differences in rodent decision making studies that may point to differences in the way that male and female rats weigh the threat of aversive outcomes against opportunities to earn food. For instance, Pellman and colleagues (2017) arranged for a previously-safe foraging patch to unpredictably deliver foot shocks during some foraging bouts. Male rats responded by reducing their total number of visits to the patch, but lengthening the duration of their visits. In contrast, female rats simply visited the now-risky patch less frequently, and eventually lost body mass as a result. In a similar vein, Orsini and colleagues (2016) measured rats’ preferences between a large food reward that was sometimes accompanied by a shock and a smaller amount of food that carried no risk of shock, and found that female rats had a stronger preference for the smaller and safer option compared with male rats. Indeed, even a simulated predator cue (Zambetti et al., 2019) had a stronger impact on food-seeking behavior in female rats despite the absence of any actual painful outcome.

In our experiment, female rats adopted a strategy that reduced the amount of time they spent in the open-field foraging patches, thereby potentially reducing predation risk. At the same time, however, female rats were more willing to cross through the exposed central region of the open fields rather than lingering near the edges like male rats. This fact, along with the relatively modest difference in residence times between male and female rats make it difficult to argue that the sex differences we observed were fully driven by differences in threat assessment. Future work combining theoretical and experimental approaches will be critical to identifying the adaptive basis (if any) of the sex differences reported here.

### Limitations and future directions

The patch-leaving task introduced here complements other approaches to studying decision making and foraging behavior in rodents, and is particularly amenable to electrophysiological studies as the structure of the task closely follows classic designs used to investigate spatial representations in the brain (Muller & Kubie, 1987). Nevertheless, future studies using this approach could improve and expand on the work reported here. Task conditions in which the difference in food earnings for good and bad foraging strategies was larger would help rule out satisficing, the use of a “good enough” but not optimal strategy, as an explanation of overstaying. Similarly, a “closed economy”, in which subjects earned their entire allotment of food by performing the task, would test whether rats were simply not energetically challenged enough to incentivize discovering and implementing better foraging strategies. In addition, testing a wider range of food outcomes would help establish the generality of these results; for instance, there is evidence for sex differences in motivation to earn outcomes with certain combinations of nutrients (Castro et al., 2022; Maric et al., 2022). Such differences could influence foraging strategies measured on the task. Relaxing certain elements of the task structure would make the foraging scenario more realistic. For instance, foragers in the wild are not constrained to visit patches in a rigid sequence, but rather take advantage of known regularities in resource distribution and other environmental factors to plan advantageous foraging trajectories (Janmaat et al., 2006, 2014; Rudebeck & Izquierdo, 2022). A free-choice version of the patch task would afford this possibility (Hall-McMaster et al., 2021). Instantaneous repletion of patches is also unrealistic, and future experiments could examine whether rats are sensitive to the rates at which patches replenish. Finally, foraging on our task was largely passive, requiring only that rats occupied one of the two foraging patches. Consequently, interesting questions regarding search strategies rats might use to localize spatially-distributed food items, the effort and cognitive costs of search, and related issues were not addressed. Relatedly, simulating the cost of traveling between patches by imposing a delay is convenient, but clearly neglects the physical effort that characterizes real-world travel. Developing a version of the task that requires actual movement to switch between patches (using a treadmill or exercise wheel, for instance) would make an interesting comparison with the present, delay-based task. These factors could be included in future versions of the task with suitable modification.

## Acknowledgements

This work was supported by a Faculty Research Grant from the UCLA Academic Senate and Whitehall Foundation Research Grant 2021-12-045. MG was supported by the Build-PODER program (NIH RL5GM118975) at California State University, Northridge. The authors thank Jenny Chen and Desmond Anderson for assistance in developing a prototype of the behavioral apparatus. Katrina Schrode provided valuable feedback on a previous version of the manuscript.

## Author contributions

M.G. and A.M.W. designed the study. M.G. and S.G. collected the data. A.M.W., M.G., and S.G. analyzed the data. A.M.W. wrote the initial draft. A.M.W., M.G., and S.G. edited the final version.

